# Comparative genomics of ten new *Caenorhabditis* species

**DOI:** 10.1101/398446

**Authors:** Lewis Stevens, Marie-Anne Félix, Toni Beltran, Christian Braendle, Carlos Caurcel, Sarah Fausett, David Fitch, Lise Frézal, Taniya Kaur, Karin Kiontke, Matt D. Newton, Luke M. Noble, Aurélien Richaud, Matthew V. Rockman, Walter Sudhaus, Mark Blaxter

## Abstract

The nematode *Caenorhabditis elegans* has been central to the understanding of metazoan biology. However, *C. elegans* is but one species among millions and the significance of this important model organism will only be fully revealed if it is placed in a rich evolutionary context. Global sampling efforts have led to the discovery of over 50 putative species from the genus *Caenorhabditis*, many of which await formal species description. Here, we present species descriptions for ten new *Caenorhabditis* species. We also present draft genome sequences for nine of these new species, along with a transcriptome assembly for one. We exploit these whole-genome data to reconstruct the *Caenorhabditis* phylogeny and use this phylogenetic tree to dissect the evolution of morphology in the genus. We show unexpected complexity in the evolutionary history of key developmental pathway genes. The genomic data also permit large scale analysis of gene structure, which we find to be highly variable within the genus. These new species and the associated genomic resources will be essential in our attempts to understand the evolutionary origins of the *C. elegans* model.

## Introduction

*Caenorhabditis elegans* has become one of the preeminent model organisms in modern biology, but only recently have we started to understand its natural ecology and evolutionary history (Félix & Braendle 2010). Placing the specifics of *C. elegans* biology in the context of its wild ecology promises to reveal the drivers for the evolution of the particular systems analysed in the laboratory, and placing *C. elegans* in the the context of its congeners and other nematodes will reveal the constraints and long-term drivers of its evolution. A large collection of wild-caught strains of *C. elegans* is now available for comparative exploration of natural variation (Cook *et al.* 2016, 2017), and ecological interactions are being investigated through co-analysis of nematodes and microbial associates from natural systems (Schulenburg & Félix 2017). In parallel, the last decade has seen a focussed search for new species in the genus *Caenorhabditis*.

The discovery of new *Caenorhabditis* species was, for many years, hindered by a poor understanding of the natural ecology of these nematodes (Félix & Braendle 2010). Surveys of natural populations of *C. elegans* and *Caenorhabditis briggsae* revealed that, rather than being “soil nematodes”, *Caenorhabditis* species thrive in microbe-rich environments, such as rotting fruits, flowers and stems (Kiontke *et al.* 2011; Félix & Duveau 2012; Félix *et al.* 2014; Ferrari *et al.* 2017). This new understanding, combined with extensive worldwide sampling efforts, has led to the discovery of more than 50 species (Kiontke *et al.* 2011; Félix *et al.* 2014; Ferrari *et al.* 2017; MAF, LF, MVR, CB, unpublished; John Wang, Michael Ailion, Erik Andersen, Asher Cutter, pers. comm.) of *Caenorhabditis*, many of which await formal species description.

Although morphology remains fundamental to species identification and diagnosis, closely related *Caenorhabditis* species are often morphologically very similar (Sudhaus & Kiontke 2007) and many morphological characters are homoplasious within the genus (Kiontke *et al.* 2011). This has motivated the use of mating tests and comparisons of molecular sequences such as ribosomal DNA (rDNA) or internal transcribed spacer (ITS) sequences, in addition to morphology, for species diagnosis (Kiontke *et al.* 2011; Félix *et al.* 2014). These molecular sequences have also enabled the reconstruction of *Caenorhabditis* phylogeny (Kiontke *et al.* 2011). Recently, whole-genome data has been exploited for phylogenomic analysis (Slos *et al.* 2017).

Genome sequences of species closely related to *C. elegans* have furthered our understanding of *C. elegans* biology and revealed the insights into the evolutionary forces that have shaped its genome. The publication of the *C. briggsae* genome in 2003 enabled the first comparative genomics studies of *Caenorhabditis* which revealed an unusually high rate of intrachromosomal rearrangement but comparatively rare interchromosomal rearrangement (Stein *et al.* 2003). Additional genome sequences have been published from species across the genus (Mortazavi *et al.* 2010; Fierst *et al.* 2015; Slos *et al.* 2017; Kanzaki *et al.* 2018; Yin *et al.* 2018). Comparisons between genomes of hermaphroditic species such as *C. elegans* and their outcrossing relatives have exposed the genomic consequences of a switch in reproductive mode from gonochorism to autogamous hermaphroditism, including changes in overall genome structure, gene structure, and protein-coding gene content (Thomas *et al.* 2012; Fierst *et al.* 2015; Kanzaki *et al.* 2018; Yin *et al.* 2018).

Here we use mating tests, accompanied by molecular and morphological analyses, to characterise and describe ten new *Caenorhabditis* species isolated from across the world. We present draft genome sequences for nine of the ten new species, and a transcriptome assembly for one, and use the data to reconstruct the *Caenorhabditis* phylogeny. By studying morphology in context of this phylogenetic tree, we find further examples of homoplasious morphological characters in the genus. We use the genome sequences to compare gene structure across the genus and to study the evolution of genes involved in a key developmental pathway. These new species and their draft genome sequences will become an important resource for the growing number of evolutionary biologists who use *Caenorhabditis* in their research.

## Results

### Ten new species declarations

By sampling a variety of substrates in diverse geographic locations, we found ten new species of *Caenorhabditis,* most of them from rotting fruit. The species described below are ingroups compared to *Caenorhabditis monodelphis* (Slos *et al.* 2017) and therefore in the *Caenorhabditis* (Osche 1952; Dougherty 1953) clade. While initial selection of isolates for further analysis was based on morphological assignment to *Caenorhabditis*, as argued and implemented in Félix *et al.* (2014), the justification for raising species is based on a biological species concept that can be easily be implemented for these culturable nematodes. Thus, for each new isolate, we attempted crosses with previously described species in culture that have the most similar ribosomal RNA cistron internal transcribed spacer 2 (rDNA ITS2) sequence (Table 1 or Table S1). The results of the crosses are shown in Table S2. For two putative species, *Caenorhabditis parvicauda* sp. n. and *Caenorhabditis uteleia* sp. n., that also had striking morphological novelty (see below), no alignment of the rDNA ITS2 region could be obtained with default parameters in NCBI nucleotide BLAST (word size 28, gap penalty 1,2, gapcosts 0,2.5). We considered these highly divergent from any known species and did not perform mating tests.

**Table 1.**
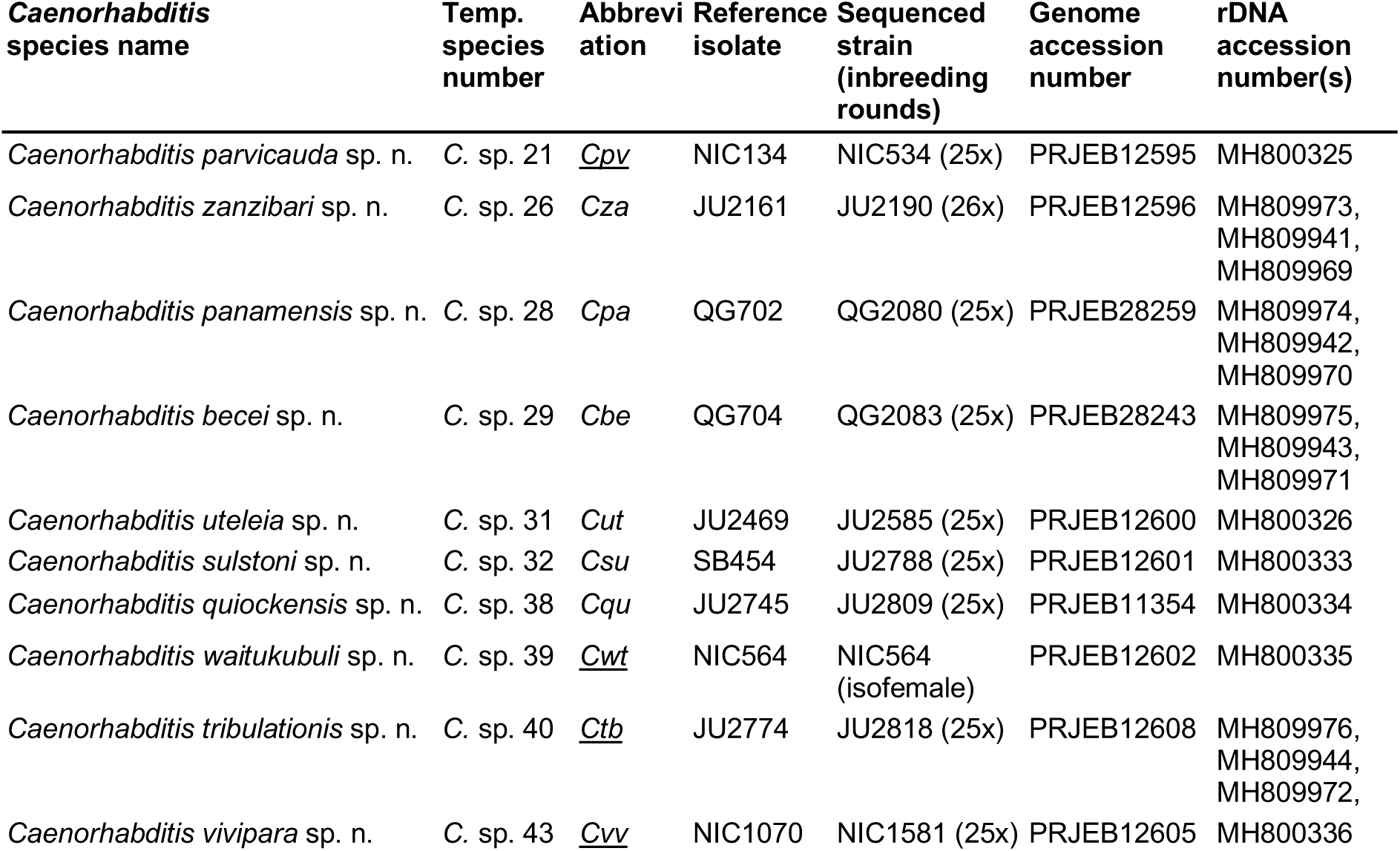
*Caenorhabditis* species names, reference isolates and sequenced strains. Specific name referents were derived as follows: 21: reduced tail (in the male). 26: first isolated in Zanzibar. 28: first isolated in Panama. 29: first isolated on Barro Colorado Island, Panama. 31: U-shape male tail. 32: in honor of John Sulston. 38: first isolated on Quiock river trail, Guadeloupe. 39: first isolated on the island of Dominique, in the local native American language. 40: first isolated in the region of Cape Tribulation, Australia. 43: viviparous.

The formal descriptions of each species can be found in Supplemental File 1. We describe the following new species (see Table 1 for the correspondence with the informal numbering system used to refer to the species in previous publications):

- *Caenorhabditis becei* sp. n.
- *Caenorhabditis panamensis* sp. n.
- *Caenorhabditis parvicauda* sp. n.
- *Caenorhabditis quiockensis* sp. n.
- *Caenorhabditis sulstoni* sp. n.
- *Caenorhabditis tribulationis* sp. n.
- *Caenorhabditis uteleia* sp. n.
- *Caenorhabditis vivipara* sp. n.
- *Caenorhabditis waitukubuli* sp. n.
- *Caenorhabditis zanzibari* sp. n.

### Morphological novelty in *Caenorhabditis parvicauda* sp. n. and *Caenorhabditis uteleia* sp. n

Morphological features of *C. parvicauda* are displayed in Fig. 1. Most striking is the absence of a fan in the tail of *C. parvicauda* adult males (Fig. 1 F,I, Fig. S5) and a left-right asymmetry in the locations of caudal papillae. The caudal papillae include 2 pairs anterior to the cloaca (v1, v2) followed by groups of 2 (v3, v4), 2 (v5 and ad) and 3 pairs (v6, v7 and pd) as seen in Fig. 1F, I and Fig. S5. Papillae ad and v5 are both open to the outside. The positions of the dorsal papillae ad (anterior dorsal) and pd (posterior dorsal) generally differ between the right and left sides at two levels. First, whereas papilla v5 has a similar anterior-posterior position on the right and left sides, papilla ad is generally anterior to v5 on the right side, and at the same position or posterior to v5 on the left side (Fig. 1F, I and Fig. S5). Second, while papilla pd has a similar anterior-posterior position on the right and left sides, ventral papillae v6 and v7 are generally located posterior to it on the right side and anterior on the left side (Fig. 1I and Fig. S5). The spicules are thick, with a complex tip (Fig. 1G). Simple pre- and post-cloacal sensilla can be seen (Fig. 1H). The males mate in a spiral position. This left-right asymmetry is highly unusual. The outer side of the mouth of *C. parvicauda* is endowed with the usual set of sensory organs disposed in a concentric manner, namely six labial sensillae, two amphids and four male-specific cephalic sensillae (Fig. 1A). The buccal cavity is short compared to most other *Caenorhabditis* (Sudhaus & Kiontke 1996), with two teeth at the base. The pharyngeal sleeve extends anteriorly to half of the buccal cavity (Fig. 1B). Three cuticular ridges can be seen in the lateral field of adults of both sexes (Fig. 1C,D). The adult female tail end is long and thin (Fig. 1E).

**Figure 1.**
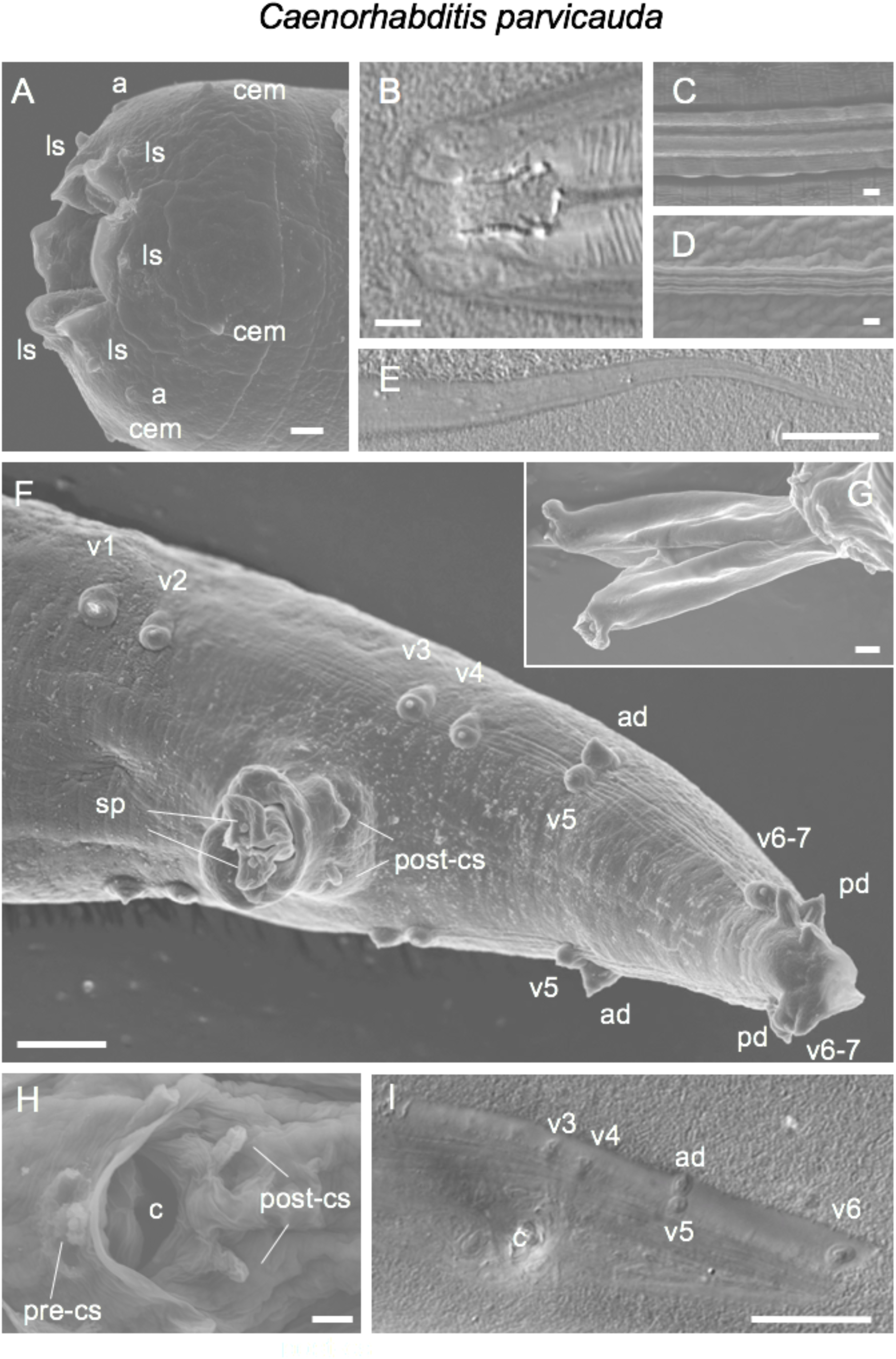
Morphology of *Caenorhabditis parvicauda* sp. n. by scanning electron microscopy (SEM) and Nomarski optics (DIC). (A,B). Mouth of an adult male (A SEM, B DIC). C. Cuticular lateral ridges of a dauer juvenile (SEM). D. Cuticular lateral ridges of an adult male (SEM). E. Female tail (DIC). F. Male tail, ventro-lateral view (SEM). G. Genital opening with extruded spicules. H. Male genital opening. I. Male tail in ventral view (DIC). Anterior is to the left in (A,B,E-I). The animals are from strain JU2070. a: amphid; ad: anterior dorsal papilla; c: cloaca; cem: male cephalic sensillum (absent in females), ls: labial sensillum; pd: posterior dorsal papilla; pre/post-cs: pre/post cloacal sensillum; v1, etc.: ventral papilla1, etc.; sp: spicule. Scale bars: 1 µm, except in (E, F, I): 5 µm. See also Fig. S5.

The morphology of *C. uteleia* is remarkable for the contour of its male tail fan (Fig. 2). The posterior margin shows a distinctive complex shape with one large central valley and two smaller ones on the left and right sides (Fig. 2C, 2D). The fan is well developed, opened anteriorly and shows a smooth, unserrated, margin. One group of two pairs of rays are found anterior to the cloaca, followed by two groups of 4 and 3 pairs. The dorsal rays are in antero-posterior positions 5 and 7. The pre-cloacal sensillum is on a hook, itself in between two characteristic lateral folds (Fig. 2C’,2D’,2F show the hook, the gubernaculum ventral end with two lateral ears and the pair of post-cloacal sensilla). The spicule tip is pointed (Fig. 2C’, 2F).

**Figure 2.**
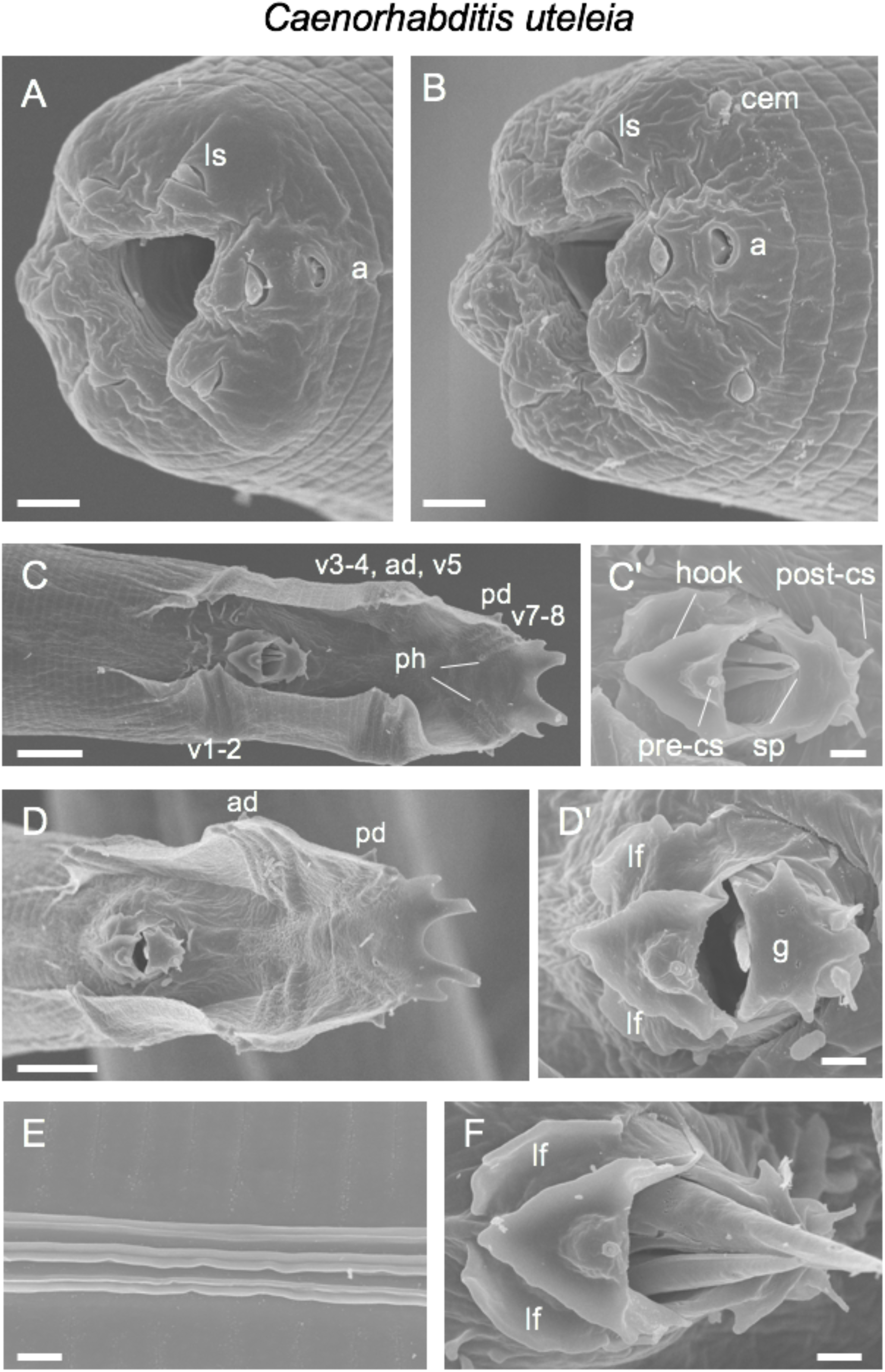
Scanning electron microscopy of *Caenorhabditis uteleia* sp. n. A. Mouth of an adult female. B. Mouth of an adult male. (C,D) Male tail in ventral view. (C’,D’) Higher magnification of the corresponding male genital openings. E. Cuticular lateral ridges, adult male. F. Male genital opening. The animals are from strain JU2469. Anterior is to the left. a: amphid; cem: male cephalic sensillum (absent in females); lf: lateral fold on either side of the hook; ls: labial sensillum; r1, etc.: ray 1, etc.; ph: phasmid; sp: spicule; pre/post-cs: pre/post cloacal sensillum; g: posterior end of the gubernaculum. Bars: 1 µm, except in (C, D): 5 µm.

### Genome sequences of nine new *Caenorhabditis* species

We sequenced the genomes of all newly-described species to high coverage (100 - 350x) using short-read Illumina technology. After identifying and removing reads from non-target organisms, we generated draft assemblies for each species, employing heterozygosity-aware assembly approaches where necessary. Draft assemblies were scaffolded using assembled transcripts or long-insert (or “mate-pair”) data, where available. The sequence data generated for *Caenorhabditis vivipara* were not of sufficient quality to generate a reference assembly and are not discussed further. All assemblies have been submitted to DDBJ/ENA/GenBank (Table 1).

Assembly span indicated substantial variation in genome size among species (Table 2). At 65.1 Mb, the genome of *Caenorhabditis sulstoni* is the smallest *Caenorhabditis* genome published thus far, and nearly 35 Mbp smaller than the *C. elegans* genome. While we note that short-read based assemblies may collapse repeats and thus yield underestimates of true span, or fail to co-assemble haploid components in very heterozygous regions and thus inflate estimates, these effects are usually minor in *Caenorhabditis* genomes (because of the low repeat content and the use of inbred strains). The contiguity of the resulting assemblies was also highly variable. The assemblies of *Caenorhabditis becei* and *C. panamensis*, which were scaffolded with long-insert data, are the most contiguous, with N50 lengths of 487 kbp and 768 kbp, respectively. The assembly of *Caenorhabditis waitukubuli* is the least contiguous, with an N50 length of 15.1 kbp. Kmer spectrum analysis (Fig. S1) indicated extensive heterozygosity present in the genome of this strain which is has not been fully collapsed during assembly. The proportion of undetermined bases (i.e., gaps denoted as Ns) was low in all cases. Despite considerable differences in assembly contiguity, BUSCO and CEGMA indicated that all assemblies were of high gene-level completeness.

**Table 2:**
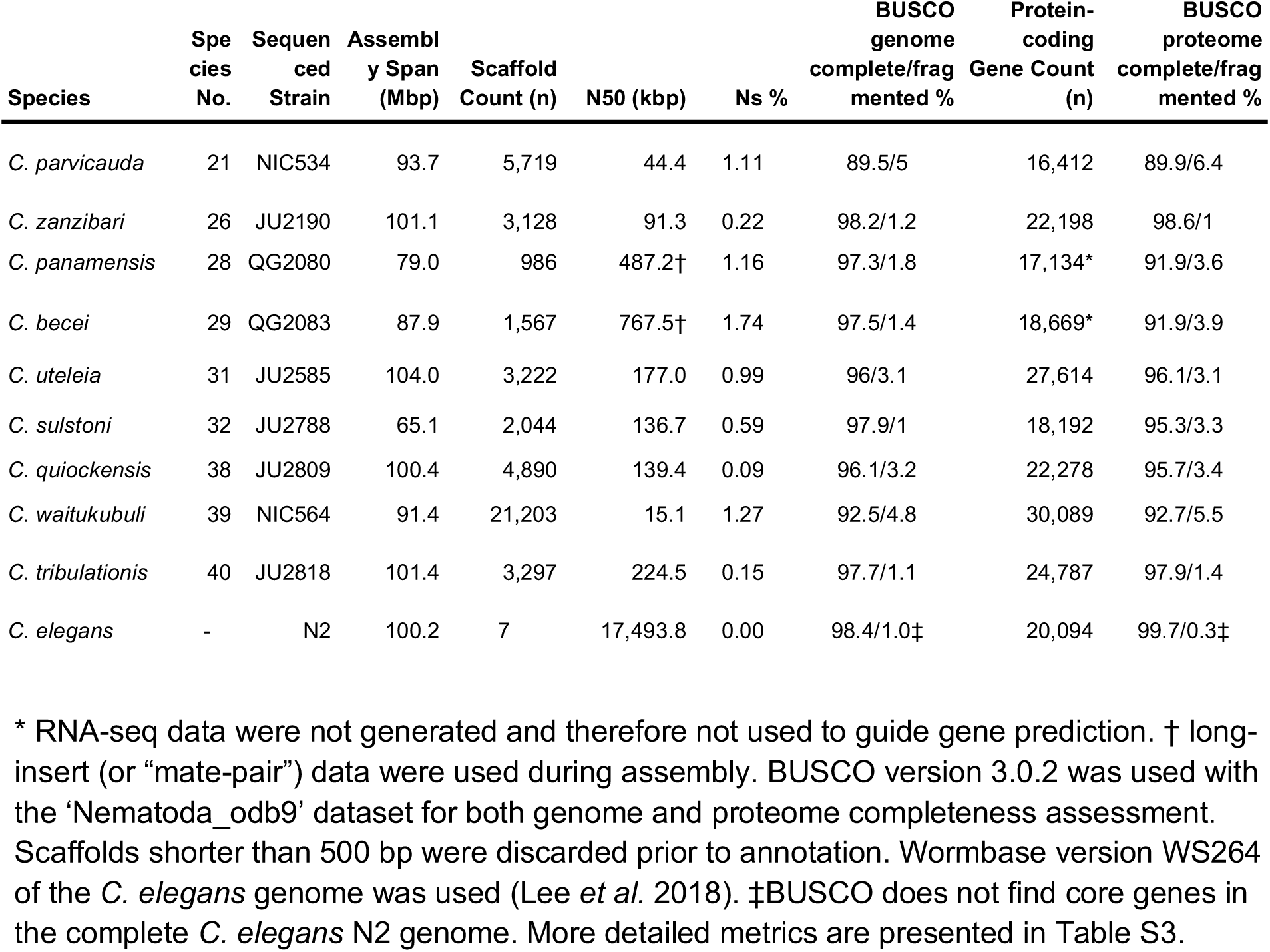
Genome assembly and annotation metrics for nine species of *Caenorhabditis*. * RNA-seq data were not generated and therefore not used to guide gene prediction. † long-insert (or “mate-pair”) data were used during assembly. BUSCO version 3.0.2 was used with the ‘Nematoda_odb9’ dataset for both genome and proteome completeness assessment. Scaffolds shorter than 500 bp were discarded prior to annotation. Wormbase version WS264 of the *C. elegans* genome was used (Lee *et al.* 2018). ‡BUSCO does not find core genes in the complete *C. elegans* N2 genome. More detailed metrics are presented in Table S3.

RNA-seq data were used to guide gene prediction for all species except *C. becei* and *C. panamensis*. The number of protein-coding genes predicted in each assembly varied considerably. The genome of *C. parvicauda* has the fewest predicted genes, at 16,412. This is likely an underestimate of the true gene number, as ∼5% of BUSCO genes were absent from the assembly. *C. waitukubuli* has 30,089 predicted protein-coding genes. 19.8% of BUSCO genes found in this assembly were present in multiple copies, suggesting that this high gene number is an assembly artefact arising from regions of uncollapsed heterozygosity present in the assembly. The lack of RNA-seq data for *C. becei* and *C. panamensis* resulted in less complete gene sets, with a larger percentage of BUSCO genes missing from the gene sets (4.5% and 4.2%, respectively) than from the draft assemblies (0.9% and 1.1%, respectively).

### Phylogenetic relationships within the genus *Caenorhabditis*

Previous analyses of *Caenorhabditis* phylogeny have used morphology or small numbers of loci and have defined subgeneric groups of taxa (Kiontke *et al.* 2011): the *Elegans* supergroup, which contains the *Japonica* and *Elegans* groups and the *Drosophilae* supergroup, which contains the *Drosophilae* and *Angaria* groups. We exploited our new and existing genomic data to reexamine the phylogenetic structure of *Caenorhabditis*. We performed orthology clustering of 781,865 protein sequences predicted from the genomes and transcriptomes of all ten newly-described species, 22 other *Caenorhabditis* species (C. elegans Sequencing Consortium 1998; Stein *et al.* 2003; Mortazavi *et al.* 2010; Kanzaki *et al.* 2018; Yin *et al.* 2018), and from the outgroup taxon *Diploscapter coronatus* (Hiraki *et al.* 2017). We identified 1,988 single-copy orthologues, each of which was present in at least 27 of the 33 taxa, and aligned their amino-acid sequences. We performed maximum likelihood (ML) and Bayesian inference (BI) analyses on a concatenated alignment of these loci. We also employed a supertree approach by estimating gene trees for all single-copy loci using ML analysis and providing the resulting topologies to ASTRAL-III (Mirarab & Warnow 2015) to estimate the species tree.

The three analyses yielded highly congruent, well supported topologies that displayed very few inconsistencies, discussed below (Fig. 3; Fig. S2-4). The majority of relationships, including the monophyly of both the *Elegans* and *Japonica* groups, were recovered with maximal support (bootstrap support of 100 and Bayesian posterior probability values of 1) regardless of method. All approaches recovered a clade of *C. guadeloupensis* + *C. uteleia* as sister to the *Elegans* supergroup. We found *C. parvicauda* to be the second-most basally arising species in the genus. We recovered *C. sulstoni, C. becei, C. waitukubuli* and *C. panamensis* as members of the *Japonica* group. The sister taxa *C. zanzibari* and *C. tribulationis* were placed as members of the *Elegans* group, being most-closely related to *C. sinica.* We recovered *C. quiockensis* as as sister to *C. angaria* + *C. castelli*.

**Figure 3:**
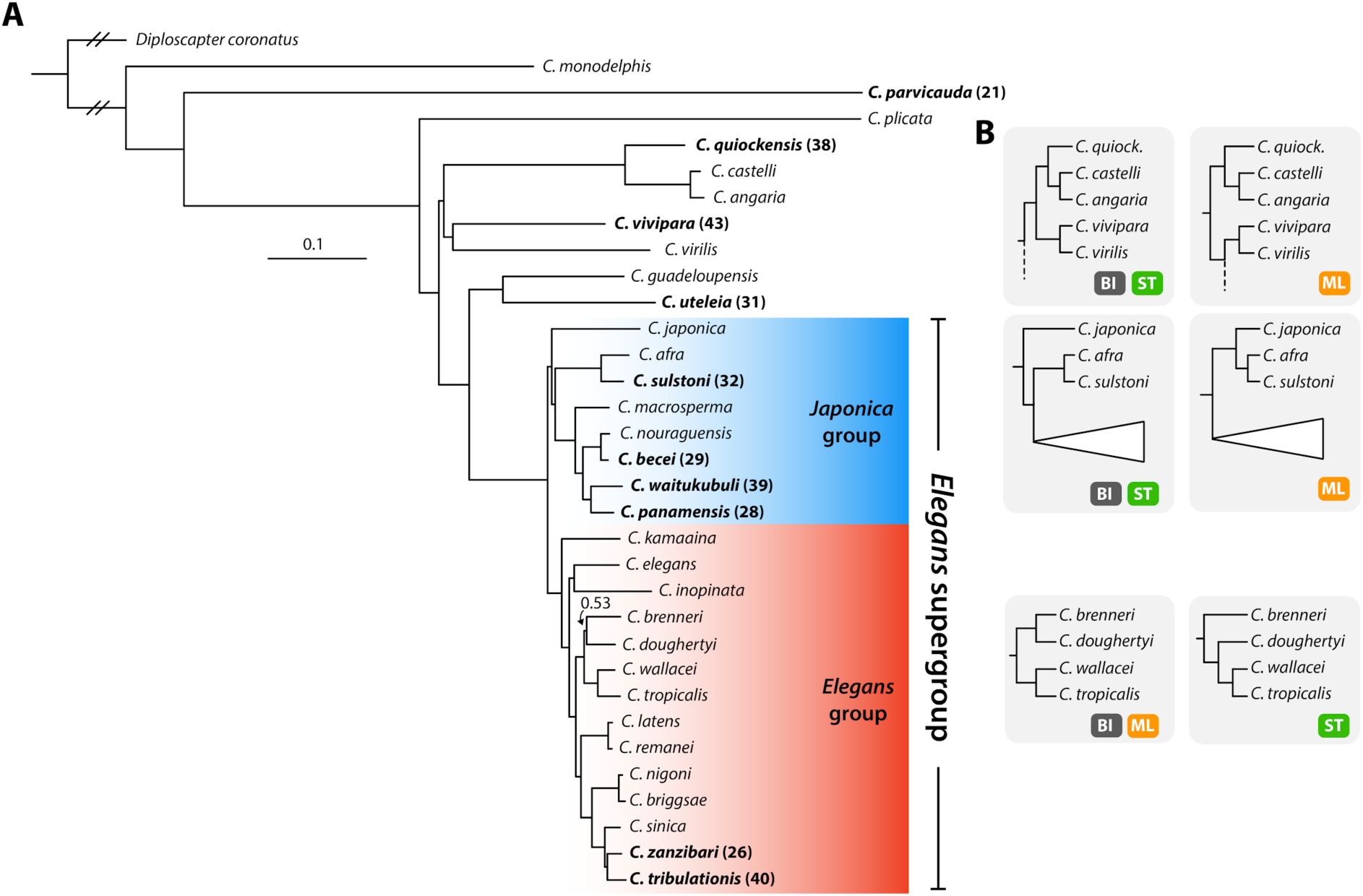
Phylogenetic relationships of 32 *Caenorhabditis* species and *D. coronatus*. **A** Phylogeny inferred using Bayesian inference with the CAT-GTR+Γ substitution model. Species described here are highlighted in bold, with previous species numbers in parentheses. Bayesian posterior probabilities are 1.0 unless noted as branch annotations. Scale is in substitutions per site. **B** Alternative hypotheses and support from each analysis approach. ML: Maximum likelihood inference using GTR+Γ substitution model, BI: Bayesian inference using the CAT-GTR+Γ substitution model, ST: Supertree approach, using gene trees as input (substitution model selected automatically for each alignment).

Three relationships, which tended to have lower-than-maximal support, were inconsistent across methods (Fig. 3B). Both the supertree approach and BI recovered the *C. vivipara* + *C. virilis* group as sister to the clade containing *C. quiockensis, C. castelli* and *C. angaria*, while ML analysis recovered this group as sister to the group consisting of the sister taxa *C. guadeloupensi*s + *C. uteleia* and the *Elegans* supergroup. Secondly, the placement of *C. japonica* as sister to other members of the *Japonica* group was recovered by both the supertree approach and by BI, whereas ML analysis recovered *C. japonica* as sister to *C. afra* + *C. sulstoni*. Lastly, BI and ML analysis recovered *C. brenneri* and *C. doughertyi* as sister taxa, whereas the supertree approach recovered *C. brenneri* as sister to the clade containing *C. doughertyi, C. wallacei* and *C. tropicalis*.

### Morphological character evolution

What are the implications of the new species and the new phylogenetic tree for character evolution in the genus? Character evolution in *Caenorhabditis* was first studied in Sudhaus & Kiontke (1996) based on a morphology-based phylogenetic tree. Kiontke *et al.* (2011) and Félix *et al.* (2014) mapped morphology onto a previous six-gene phylogenetic tree and recently Slos *et al.* (2017) discussed the stem species pattern using a morphological comparison with the sister group Protoscapter.

Starting from the base of the genus, defined by the placement of the sister taxon Protoscapter (Slos *et al.* 2017), containing *Diploscapter*, we now have as successive branches *C. monodelphis* (likely to be associated with the presently unavailable *Caenorhabditis sonorae*)(Slos *et al.* 2017), then *C. parvicauda* and then *C. plicata*. Species in these three branches share characters that are ancestral in the genus (plesiomorphous), such as an absence of hook. All four species differ greatly in morphology and show private characters and combinations thereof. *C. parvicauda* is unusual for *Caenorhabditis* because the male tail does not include an extended fan and papilla v5 does not appear broader than the others. However, in contrast to *C. monodelphis, C. parvicauda* displays some characters that are shared with other *Caenorhabditis* species, such as three lateral cuticular ridges. *C. parvicauda* also shares some characters with some other species that are likely homoplastic. For example, *C. parvicauda* mate in a spiral fashion on an agar plate, as do species of the *Angaria* group.

*C. quiockensis* is typical for the *Angaria* group (*C. angaria, C. castelli*) (Kiontke *et al.* 2011; Félix *et al.* 2014) in displaying spiral mating and having a distinctive male tail morphology (Fig. S6A-C). The fan has an oval shape in ventral view and is open anteriorly. Rays 4 (=ad) and 7 (=pd) open dorsally to the velum.

*C. vivipara* is distinct in its viviparity. In the closely related species *C. virilis*, females lay late-stage embryos compared to most *Caenorhabditis* species (which accumulate few embryos in their uterus) but is not viviparous in standard laboratory conditions. Like *C. virilis* however, *C. vivipara* has a wide and heart-shaped male fan, similar to the male fans of species in the *Elegans* supergroup yet with no terminal notch (compare Fig. S6E and Fig. S7).

*C. uteleia* has a previously unknown combination of characters that do not all match its most closely related known species *C. guadeloupensis* (Kiontke *et al.* 2011; Félix *et al.* 2014). For example, the male fan of *C. uteleia* is open and unserrated on the anterior side as observed in many species in the *Drosophilae* supergroup and in basally branching species. Conversely, the anterior-dorsal (ad) ray is in the fifth position in *Elegans* supergroup species and basally branching species such as *C. monodelphis* (Slos *et al.* 2017) but not in *C. guadeloupensis.* The latter, like *C. plicata, C. drosophilae,* the Angaria group, *C. virilis* and *C. vivipara* show a dorsal ray in antero-posterior position 4 (Kiontke *et al.* 2011; present data). Thus, these characters remain homoplastic in the new phylogenetic tree and will most likely remain so. Finally, the new clade that includes *C. guadeloupensis, C. uteleia*, and the *Japonica + Elegans* groups is distinguished by the presence of a hook bearing the pre-cloacal sensillum on the anterior margin of the cloaca. This character also appears in *C. portoensis* but not in *C. virilis* (Kiontke *et al.* 2011; Félix *et al.* 2014) nor *C. vivipara* (Fig. S6F,G), and thus it will be important to affirm the position of *C. portoensis* (currently resolved in a clade including *C. virilis* (Kiontke *et al.* 2011; Félix *et al.* 2014)) to define how many times this character has evolved.

In contrast, the clade including the *Elegans* and *Japonica* groups displays little variation except for the tropical fig species *C. inopinata* (Kanzaki *et al.* 2018). One character variation that is unique to the *Elegans* group is the independent evolution of hermaphroditism in three species (Kiontke *et al.* 2011; Félix *et al.* 2014). The present work does not add new hermaphroditic species, as all ten new species reproduce through females and males. However, using the large set of species studied here including six new species in the *Elegans* + *Japonica* groups, we distinguished and mapped on the phylogenetic tree three characters in the male tail that vary in either the *Elegans* or *Japonica* group (i.e. hook shape, ray 4 length relative to adjacent rays, shape of the spicule ventral tip).

Within the *Elegans* group the small clade containing *C. sinica, C. zanzibari* and *C. tribulationis* displays a characteristic shape of the male hook with three marked lobes (Fig. 4). The hook of other species of the *Elegans* supergroup such as *C. elegans* displays a single lobe. Based on parsimony, the complex shape of the hook as first described for *C. sinica* as a specific character (Huang *et al.* 2014) now constitutes a clear apomorphy (derived character), with no pattern of convergent evolution.

**Figure 4.**
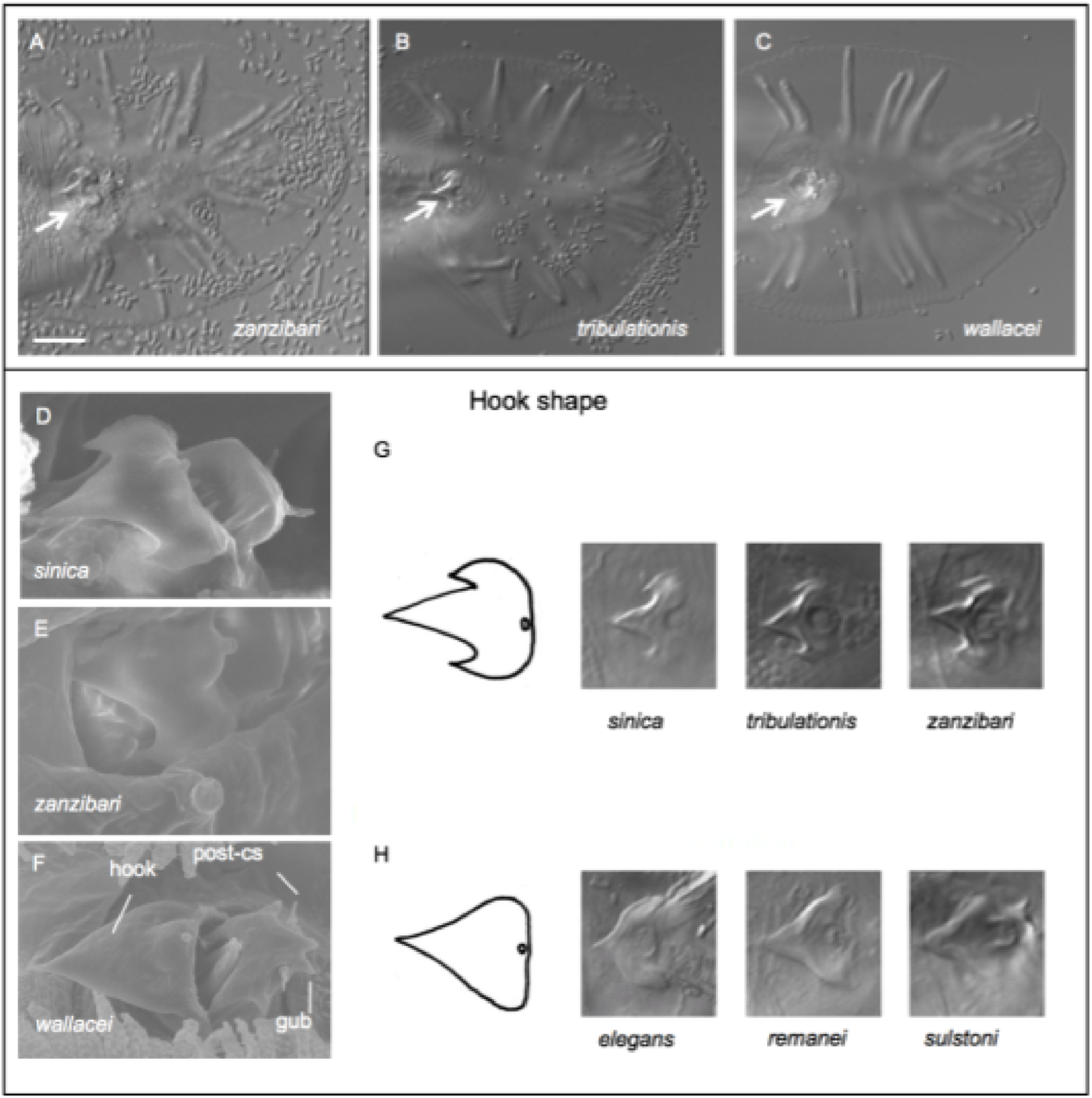
Male hook shape in *Elegans* group species. **A-C:** Ventral views of the male tail of *Caenorhabditis zanzibari* strain JU2161 (**A**), *Caenorhabditis tribulationis* strain JU2774 **(B)**, and *Caenorhabditis wallacei* JU1873 **(C),** in Nomarski optics. The arrow points to the precloacal hook. Bar: 10 µm. **D-F:** Scanning electron micrographs of the hooks and post-cloacal sensilla (post-cs) in *Caenorhabditis sinica* JU727 **(D)**, *C. zanzibari* JU2161 **(E)** and *C. wallacei* JU1873 **(F)**. “gub”: forked posterior end of gubernaculum. **G,H**: Drawings of hook shape. **(G)** refers to the trilobed shape in *C. sinica, C. tribulationis* and *C. zanzibari* sp. n, while **(H)** refers to the simpler shape in most other Elegans supergroup species such as *C. elegans, C. remanei* and *C. sulstoni*.

In the *Japonica* group, ray v4 is shorter in the clade including *C. becei, C. waitukubuli, C. panamensis*, and *C. nouraguensis*, compared to *C. sulstoni, C. afra, C. elegans* and many other species where ray v4 displays a similar length to the ad (anterior dorsal) ray (see also drawings in Kiontke et al. (2011)). Ray v4 is also short in *C. japonica* (Kiontke *et al.* 2002), but longer in *C. macrosperma* JU1857, which implies events of homoplasy in any of the configurations of the phylogenetic tree for this latter species (Fig. 3).

Kiontke *et al.* (2011) and noted that the ventral tip of the spicules was broad and complex outside the *Elegans* supergroup as well as for *C. japonica* and *C. afra*. The ventral tip of the spicules is wide in *C. afra* as well as its sister species *C. sulstoni* (Fig. S8). It also seems broader and bent at an angle in *C. becei*, more so than in *C. panamensis* or *C. waitukubuli* (Fig. S8). This character thus varies in the *Japonica* group, while the tip remains pointed in the *Elegans* group, and given the present phylogeny, some homoplasy must also be present in this character.

### Intron abundance is highly variable within *Caenorhabditis*

With the resolved phylogeny and the now extensive genome data, we have been able to initiate analysis of genus-wide processes of genome evolution in *Caenorhabditis*. Gene structure is known to vary between related *Caenorhabditis* species (Kent & Zahler 2000; Stein *et al.* 2003; Slos *et al.* 2017). Several studies using individual or small numbers of protein-coding loci have indicated that there has been a high rate of intron turnover in the genus, with intron loss being more common than intron gain (Cho *et al.* 2004; Kiontke *et al.* 2004). We studied intron abundance in the genomes of 27 *Caenorhabditis* species and *D. coronatus*.

We identified 990 single-copy orthologues present in all 28 species and counted the number of introns in each gene. We found that intron abundance was highly variable across the phylogeny, with basal species and the outgroup taxon *D. coronatus* having substantially more introns per gene than ingroup taxa (Fig. 5A). For example, a total of 10,446 introns were present in the 990 loci in *C. monodelphis* (an average of 10.5 per gene), while 6,278 introns were present in *C. elegans* (an average of 6.3 per gene).

**Figure 5:**
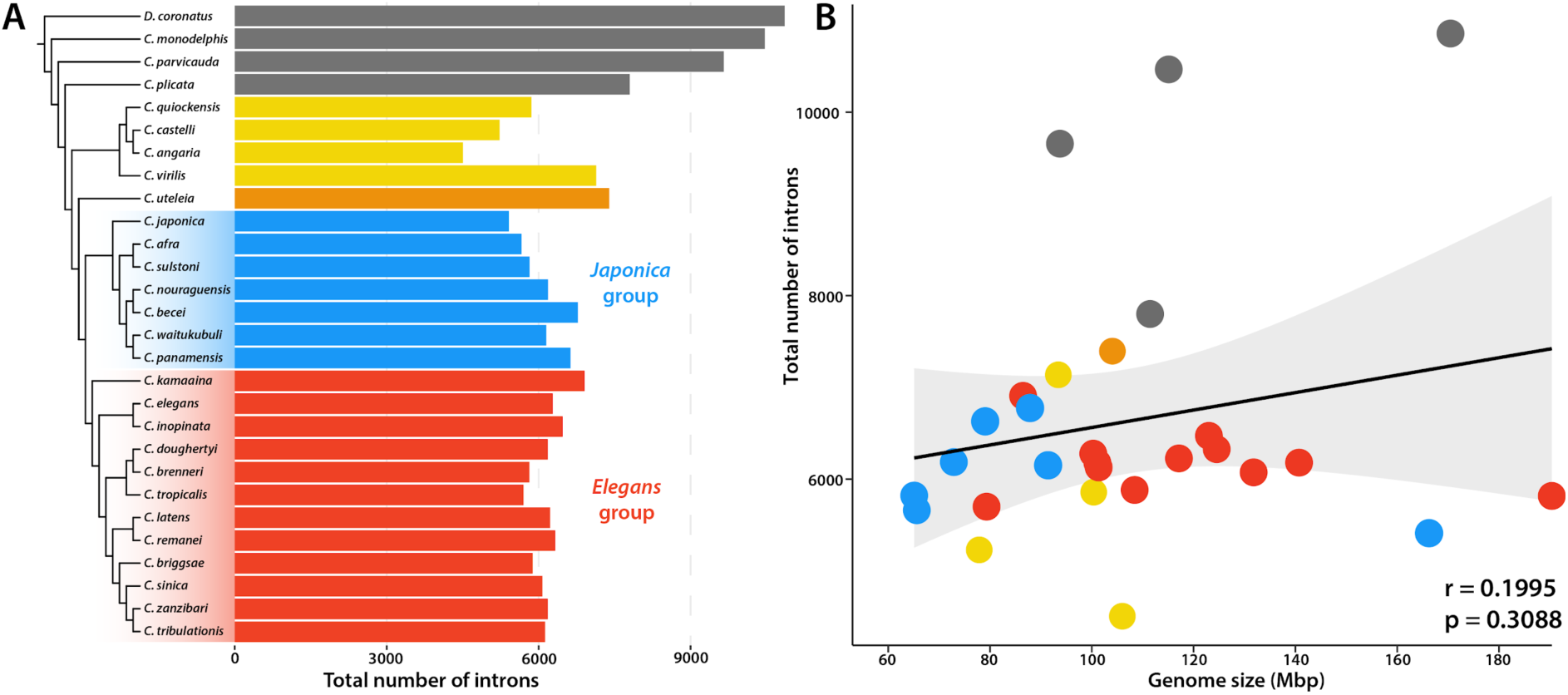
Intron number in 27 species of *Caenorhabditis* and the outgroup species *D. coronatus*. **A:** Total number of introns present in 990 single copy orthologues. The phylogenetic tree is based on Fig. 3A, with major clades highlighted. **B:** Genome size is not significantly correlated with total intron count in 990 single-copy orthologues (r=0.1995081, n=28, p=0.308). Pearson’s r analysis was performed in R.

Surprisingly, we found that intron abundance was not significantly correlated with genome size (as indicated by assembly span) in *Caenorhabditis* (Fig. 5B). For example, the genomes of *C. afra* and *C. sulstoni* are both 35% smaller than the genome of *C. elegans*, but have only 7 and 10% fewer introns, respectively. In contrast, the genome of *C. monodelphis*, the most basal *Caenorhabditis* species, has 67% more introns than does *C. elegans*, while being only 15% larger.

### The evolution of Notch/LIN-12 signalling proteins in *Caenorhabditis*

The new genome data and the resolved phylogeny permits detailed examination of the origins and diversification of genes and gene families. Developmental genetics in *C. elegans* has identified many genetic systems critical across Metazoa, but has also identified idiosyncrasies that are not shared. As an example of the power of these new data, we examine Notch signalling.

Notch signalling is a highly conserved intercellular signalling pathway involved in an array of cell fate decisions during animal development (Artavanis-Tsakonas *et al.* 1999). Basic components of this pathway were characterised in *C. elegans* at the same time as in *Drosophila melanogaster* (Greenwald 1985; Greenwald 2005). Central to this pathway are the Notch receptors (Greenwald 1985; Yochem *et al.* 1988; Yochem & Greenwald 1989). These transmembrane proteins bind extracellular ligands of the Delta/Serrate/LAG-2 (DSL) family. Ligand binding results in cleavage and nuclear translocation of an intracellular domain which associates with transcription factors to influence the expression of genes involved in cell-fate decisions (Greenwald 2005). *C. elegans* possesses two Notch-like receptor loci, *lin-12* and *glp-1*, which have overlapping but not identical biological roles (Lambie & Kimble 1991). The two loci are the product of a gene duplication event, which has been followed by some degree of subfunctionalization (Rudel & Kimble 2002). We used the new genomic data to study the evolution of these two loci.

From a orthology clustering analysis including 27 *Caenorhabditis* species and the outgroup taxon *Diploscapter coronatus,* we identified the orthogroup containing the *C. elegans* proteins LIN-12 and GLP-1. We aligned the protein sequences of each member of the orthogroup and reconstructed a phylogenetic tree using maximum likelihood inference. The resulting topology (Fig. 6A) reaffirms that *lin-12* and *glp-1* are the product of a gene duplication, and indicates that this occurred at the base of the *Elegans* supergroup. The genomes of several species outside the *Elegans* supergroup, including the basal taxa *C. monodelphis* and *C. parvicauda,* encode only a single Notch-like receptor. We also find evidence for a second duplication of *glp-1* at the base of the *Elegans* supergroup, followed by loss in species belonging to the *Elegans* group, including *C. elegans* (Fig. 6B). Genomes of species belonging to the *Japonica* group have retained both copies of the *glp-1*-like gene, and therefore encode three Notch-like receptor genes in total. Both duplication branches had maximal bootstrap support. Further, within-species duplications of the Notch-like receptor genes appear to have occured in several species, including in *C. uteleia.* In those cases where the subtending branches are extremely short, such as in *D. coronatus*, these putative within-species duplications are likely to be an artefact arising from regions of uncollapsed heterozygosity present in the genome assembly.

**Figure 6:**
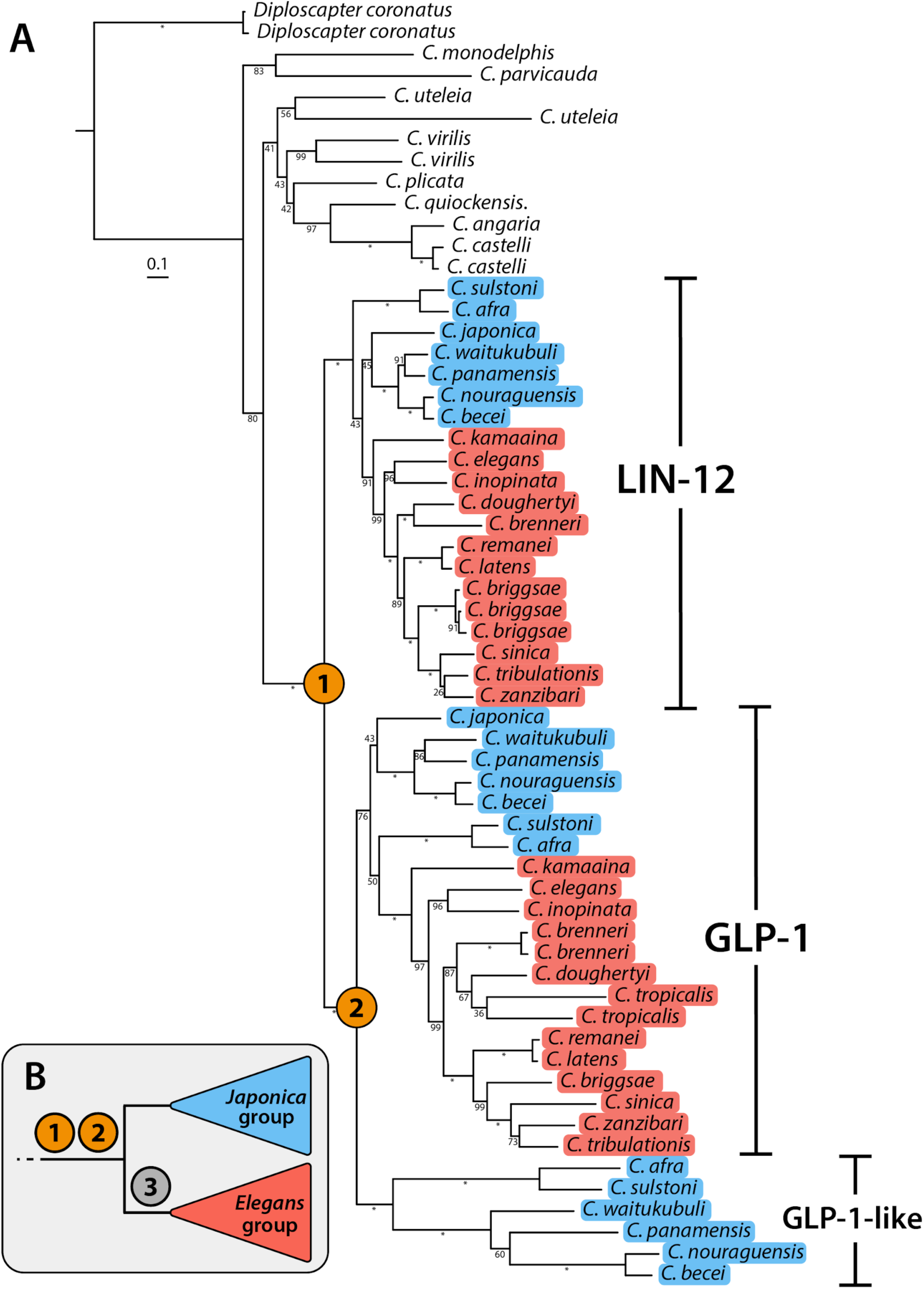
Maximum likelihood gene tree of Notch-like receptors in *Caenorhabditis.* **A:** Gene tree of the orthogroup containing *C. elegans* proteins LIN-12 (CELEG.R107.8) and GLP-1 (CELEG.F02A9.6) inferred using maximum likelihood (JCCMut+Γ substitution model). *Elegans* and *Japonica* groups are highlighted in red and blue, respectively. Duplication events are denoted by orange circles. Branch lengths represent the number of substitutions per sites; scale is shown. Bootstrap values are displayed as branch annotations, * = 100. **B:** Inferred events: 1. Duplication of ancestral Notch-like receptor gene; 2. Duplication of *glp-1* gene; 3. Loss of one of the two duplicated *glp-1* genes in the *Elegans* group.

Both *Elegans* supergroup duplication events were followed by divergence in the rate of substitution of the paralogues, as indicated by branch lengths (Fig. 6A). GLP-1 and its orthologues underwent increased rates of substitution after the first duplication event relative to LIN-12 and orthologues, with means of 0.93 and 0.58 substitutions per amino acid site, respectively (Table S4). The paralogues of GLP-1 found in the *Japonica* group have also undergone increased rates of substitution relative to GLP-1 after the second duplication event, with means of 1.22 and 0.78 substitutions per site, respectively.

The binding of DSL ligands by Notch receptor proteins is mediated by extracellular epidermal growth factor (EGF) repeats (Rebay *et al.* 1991). In *C. elegans*, LIN-12 and GLP-1 differ in the number of EGF repeats, with 13 and 10 respectively. To infer the structure of the ancestral *Caenorhabditis* Notch-like receptor, we counted the number of EGF repeats in all orthologues of LIN-12 and GLP-1 using InterProScan. The single Notch-like receptors of those species that do not belong to the *Elegans* supergroup usually possess 13 EGF repeats (Table S5), suggesting the EGF-repeat structure of the ancestral Notch-like receptor was more similar to that of LIN-12 than to that of GLP-1.

## Discussion

### Phylogenetic relationships in the genus *Caenorhabditis*

Using genomic and transcriptomic data from 32 species of *Caenorhabditis* and the outgroup taxon *D. coronatus*, we have generated the most comprehensive *Caenorhabditis* phylogenetic tree published to date, in terms of number of taxa and number of orthologues sampled. The majority of relationships were recovered with maximal support by all analyses and are consistent with previous studies (Kiontke *et al.* 2011; Slos *et al.* 2017). The ten species described here are sampled from all previously-defined major clades in the genus, and include the second-most early diverging *Caenorhabditis, C. parvicauda.*

Our results corroborate the monophyly of the *Japonica* and *Elegans* groups (and therefore the *Elegans* supergroup) as defined in Kiontke *et al.* (2011). The majority of relationships within these groups are consistent with previous studies (Kiontke *et al.* 2011; Slos *et al.* 2017). However, *C. kamaaina* (sp. 15), which in our analyses was recovered as the most early diverging species in the *Elegans* group, has previously been recovered as sister to the *Japonica* group (Kiontke *et al.* 2011). Our finding that the *C. guadeloupensis* + *C. uteleia* group is sister to the *Elegans* supergroup is consistent with the tree topology in Slos *et al.* (2017) and provides evidence that the *Drosophilae* supergroup, as recovered by the analysis of Kiontke *et al.* (2011), is paraphyletic.

Interestingly, relationships at three nodes were inconsistent across analyses. The placement of *C. japonica* as sister to *C. afra* + *C. sulstoni* and the placement of the clade containing *C. vivipara* + *C. virilis* as sister to the *Elegans* supergroup were recovered by ML analysis only. Both of these relationships disagree with all previously published *Caenorhabditis* phylogenetic trees (Kiontke *et al.* 2011; Slos *et al.* 2017). The sister relationship of *C. brenneri* and *C. doughertyi,* as recovered by BI and ML, was recovered in the analyses of Kiontke *et al.* (2011), but not by Slos *et al.* (2017).

It is not clear why the different approaches generate inconsistent topologies, and whether these inconsistencies are due to conflict in the data or due to low resolution which result in ambiguous regions of the topology. Concatenated alignments of large numbers of loci, such as the one used in our BI and ML analyses, may contain genes which have different histories due to processes such as incomplete lineage sorting or introgression through hybridisation in the early stages of speciation (Maddison & Wiens 1997). Given sufficient conflicting signals, concatenation approaches are known to lead to inaccurate topologies (Kubatko & Degnan 2007). Coalescent-based summary approaches, such as ASTRAL-III, are able to accommodate conflicting signals in gene trees arising from incomplete lineage sorting (Mirarab *et al.* 2014), but may be sensitive to gene tree estimation error (Gatesy & Springer 2014; Roch & Warnow 2015). The different models used in BI and ML analysis (CAT-GTR and GTR, respectively) are presumably one source of conflict between these two analyses. The CAT-GTR model, unlike the GTR model, is able to accomodate heterogeneity in the amino acid substitution process at different sites in the alignment (Lartillot & Philippe 2004). Further work is required to determine whether accounting for such heterogeneity is necessary for model adequacy for this dataset.

We feel it premature to redefine the groups and supergroups in *Caenorhabditis* until we have more complete representation of the known species, and particularly of species currently placed in the *Drosophilae* group. Our data could be used to support a broadening of the *Elegans* supergroup to include *C. uteleia* and *C. guadeloupensis*. Alternately, the *Elegans* supergroup could be retained as the clade including the last common ancestor of *C. elegans* and *C. japonica* and their respective groups, but not *C. uteleia* and *C. guadeloupensis*. Many more species outside the *Elegans* and *Japonica* groups are now available in culture for genome sequencing and subsequent placement on the phylogenetic tree (Kiontke *et al.* 2011; Félix *et al.* 2014). As sequencing technologies improve, and chromosomal assembly becomes more achievable (Kanzaki *et al.* 2018) we would also hope to use a chromosomally-resolved genomic tree to uncover additional patterns and reveal novel processes in *Caenorhabditis* evolution.

### Intron abundance is highly variable within *Caenorhabditis*

By comparing the structure of 990 single-copy loci present in the genomes of 27 *Caenorhabditis* species and the outgroup taxon *D. coronatus,* we found a striking pattern of variability of intron abundance, with basal taxa having substantially more introns per gene than ingroup taxa. This finding is consistent with previous studies of gene structure in *Caenorhabditis* which have used much smaller numbers of loci or species (Kent & Zahler 2000; Stein *et al.* 2003; Cho *et al.* 2004; Kiontke *et al.* 2004; Slos *et al.* 2017).

The pattern of variation in intron abundance in the genus could have arisen as a result of a bias towards intron gain in basal and outgroup taxa or due to an ongoing bias towards into loss in the ingroup taxa. Previous analyses of small numbers of protein-coding genes revealed intron loss to be more common than intron gain in *Caenorhabditis* (Robertson 1998; Cho *et al.* 2004; Kiontke *et al.* 2011). Indeed, many reported instances of intron gain in *C. elegans* and *C. briggsae* (Coghlan & Wolfe 2004) were subsequently revealed to be the product of multiple independent losses when more taxa were included (Roy & Penny 2006). Therefore, we believe it is likely that this pattern is the product of extensive intron loss in ingroup taxa and that *C. monodelphis* represents an intron-rich ancestral state.

The mechanisms of intron gain and loss, and the evolutionary forces that govern their rates, remain unclear (Fedorov *et al.* 2003; Roy & Gilbert 2005; Roy & Irimia 2009). Changes in intron number could contribute to differences in overall genome size. However, we found that intron abundance was not significantly correlated with genome size, suggesting that the evolutionary forces that govern rates of intron loss and gain are distinct from those which govern genome size expansion and contraction. Future, more detailed studies of intron gain and loss in *Caenorhabditis* may reveal clues as to which forces govern their rates and, importantly, which mechanisms are responsible for these events. Genome sequencing projects for several other *Caenorhabditis* species are currently underway, and the resulting data will be vital to these efforts.

### The evolution of Notch/LIN-12 signalling proteins in *Caenorhabditis*

*C. elegans* is a major developmental biology model, and understanding how its particular instance of nematode (and animal) development was established may reveal the logic of the evolution of development and the importance of processes such as developmental systems drift. The availability of complete genomic information for species in the genus *Caenorhabditis* will facilitate realisation of this research programme. We studied the evolution of Notch-like receptor genes in the genomes of 27 species of *Caenorhabditis* and in *D. coronatus.* We identified that the *C. elegans* genes *lin-12* and *glp-1* are the product of a gene duplication event that occured at the base of the *Elegans* supergroup. We also found evidence of a subsequent duplication of *glp-1* on the same branch, followed by loss at the base of the *Elegans* group. The genomes of *Japonica* group species have three Notch-like receptor genes. We found that the *lin-12* and its orthologues were more conserved, with a lower rate of substitution compared to the loci present in more basal species than the *glp-1* orthogroup members and an EGF-repeat structure that is more similar to that of the inferred ancestral Notch-like receptor.

Previous genetic analyses of *lin-12* and *glp-1* identified several shared and several receptor-specific developmental functions, suggesting subdivision of ancestral roles after duplication (Lambie & Kimble 1991). While LIN-12 and GLP-1 proteins have diverged in sequence, it is likely that a substantial amount of this divergence in function is due to difference in expression patterns of these two proteins as GLP-1 is capable of performing LIN-12-specific roles when placed under the control of *lin-12* regulatory sequences (Fitzgerald *et al.* 1993). The extent to which ancestral roles have been subdivided (subfunctionalisation), or whether novel roles have been acquired (neofunctionalisation), and to what extent these changes are due to changes in expression pattern, could be addressed by studying the function of the ancestral Notch-like receptor gene in species outside the *Elegans* supergroup. Particularly exciting are the third Notch-like receptor genes present in the genome of *Japonica* group species which appear to have undergone a substantial increase in substitution relative to *glp-1* and its orthologues after duplication. Genetic analyses of Notch-like receptors, which have yet to be carried out in these species, could reveal the implications of this increased rate of substitution, including whether further subdivision of ancestral roles has occurred or novel roles have been acquired.

### Genomics, species description and *Caenorhabditis* biology

By describing new species of *Caenorhabditis* with high quality genomic data we hope to promote not only understanding of the evolution of this exciting genus, but also exploitation of these new species in deepening understanding of the pattern and process of organismal evolution. In particular, the ability to interfere with gene function through RNAi and accurately modify genomes using CRISPR-Cas editing is routine in *C. elegans* and becoming established in other species. Using the genomic data presented here, specific loci can be deleted or altered to test hypotheses of gene function in a rich phylogenetic context.

Currently, strains corresponding to about 50 species of *Caenorhabditis* are available in culture. Our goal is to provide genome sequences and formal species descriptions for all these species. The assemblies presented here are not chromosomal, unlike the genome of *C. elegans*, because of the inability of short-read data to bridge and resolve large repeats. Recent developments in genomic technologies make phased, chromosomal assemblies of all *Caenorhabditis* species’ genomes an achievable goal, and we and others are using long read and other approaches to advance this goal.

Approximately 20,000 new species are described each year. We also suggest that, wherever technically possible, the generation of genomic data as part of the introduction of new taxa should become standard. Complete sequencing of all taxa on earth has been proposed (Lewin *et al.* 2018), and active support should be offered to taxonomists to add their new species to this global effort.

## Methods

### Sampling and isolation

See Kiontke et al. (2011) and Barrière & Félix (2014) for details of sampling strategies. Briefly, rotting vegetable matter samples were collected and stored in plastic bags. The samples were then analyzed in the laboratory by placing them onto 90 mm NGM agar plates (Stiernagle 2006), seeded in the centre with *Escherichia coli* OP50. *Caenorhabditis* species are generally attracted by the *E. coli* lawn. Isofemale lines were established by isolation of a single female that was already mated or by co-culturing one female and one male. A list of isolates is available as Table S1.

### Culture and freezing

Nematodes were cultured on NGM agar plates and frozen with standard *C. elegans* protocols (Stiernagle 2006). Some species, such as *C. parvicauda* do not grow well on *E. coli* OP50 and are better kept with some of their original microbial environment. Like other species in the *Angaria* group, *C. quiockensis* does not survive well freezing and thawing with the standard *C. elegans* protocol and was heat-shocked for 1-2 hrs at 37°C before freezing.

### Crosses

5 L4 females and 5 L4 or adult males were placed together on a 55 mm agar plate seeded with *E. coli* OP50. The plate was checked regularly for presence and cross-fertility of the progeny.

### Inbreeding and nucleic acid preparation

For inbreeding, one L4 female and one male were used to seed each generation. Inbreeding was performed for 20-25 generations.

After thawing, each inbred strain was bleached and grown on 90 mm NGM plates enriched with agarose (for 1 L: 3 g NaCl, 5 g bacto-peptone, 10 g agar, 7 g agarose, 1 mL cholesterol 5mg/mL in ethanol, 1ml CaCl_2_ 1M, 1 mL MgSO_4_ 1 M, 25 mL KPO_4_ 1 M). Worms were harvested just after starvation and washed in M9 several times to remove *E. coli.*

For genomic DNA extraction, the nematode pellets were resuspended in 600 µL of Cell Lysis Solution (Qiagen) complemented with 5 µl of proteinase K (20 µg/µl) and incubated overnight at 56°C with shaking. The day after, the lysates were incubated one hour at 37°C with 10 µL of RNAse A (20 µg/µL) and the proteins were precipitated with 200 µL of protein precipitation solution (Qiagen). After centrifugation, the supernatants were collected in new tubes and genomic DNA were precipitated with 600 µL of isopropanol. The pellets were washed in ethanol 70% and dried one hour before being resuspended in 50 µL of DNAse free-water.

For RNA extraction, 100 µL of nematode pellet was resuspended in 500 µL of Trizol (5 volumes of Trizol per volume of pelleted nematodes). The Trizol suspension was frozen in liquid nitrogen and then transferred to a 37°C water bath to be thawed completely. This freezing/thawing process has been repeated 4-5 times and the suspension was vortexed 30 s and let rest 30 s (5 cycles). 100 µL of chloroform was added and the tubes were shaken vigorously by hand for 15 s and incubated 2-3 min at room temperature. After centrifugation (15 min at 13000 rpm and 4°C), the aqueous (upper) phase containing the RNA was transferred to a new tube and precipitated with 250 µL of isopropanol. The pellets were washed in 70% ethanol and dried 15-20 min before being resuspended with 50-100 µL of RNAse-free water. An aliquot of each DNA and RNA preparation was run on agarose gel to check their quality and quantitated with Qubit (Thermo Scientific)

### Genome sequencing

For *C. parvicauda, C. quiockensis, C. sulstoni, C. tribulationis, C. uteleia,C. waitukubuli* and *C. zanzibari,* two short-insert (insert sizes of 300 bp and 600 bp, respectively) genomic libraries and a single short-insert (150 bp) RNA library were prepared using Illumina Nextera reagents and sequenced (125 bases, paired-end) on an Illumina HiSeq 2500 at Edinburgh Genomics (Edinburgh, UK). For *C. becei and C. panamensis* a short-insert (insert size of 600 bp) genomic library and a long-insert (insert size of 4 kb) genomic library were prepared using Illumina Nextera reagents and sequenced (100 bases, paired-end) on an Illumina HiSeq 2500 at the New York Genome Center (New York, USA). All raw data have been desposited in the relevant International Nucleotide Sequence Database Collaboration (INSDC) databases (Table 1).

### *De novo* genome assembly and gene prediction

Detailed methods for each species, along with all software tools used (including versioning and command line options), are available in Supplementary File 2. We performed quality control of all raw sequence data using FastQC (Andrews & Others 2010) and used Skewer (Jiang *et al.* 2014) and FASTQX Toolkit (Gordon & Hannon 2010) to remove low-quality bases and adapter sequence. Adapters were removed from long-insert data using NextClip (Leggett *et al.* 2014). For each species, we identified contaminants using taxon-annotated, GC-coverage plots as implemented in blobtools (Laetsch & Blaxter 2017) (preliminary assemblies were generated using CLC Bio (CLCBio, Copenhagen, Denmark), and likely taxon origin determined using NCBI-BLAST+ (Camacho *et al.* 2009) or using Kraken (Wood & Salzberg 2014). Reads originating from contaminant genomes were discarded. We estimated the optimal k-mer length for assembly independently for each genome using KmerGenie (Chikhi & Medvedev 2014). Preliminary assemblies were generated using several de Bruijn graph assemblers, including Velvet (Zerbino & Birney 2008) and Platanus (Kajitani *et al.* 2014), across several parameter values. The resulting assemblies were assessed using numerical metrics, and two biological completeness metrics, Core Eukaryotic Genes Mapping Approach (CEGMA) (Parra *et al.* 2007) and Benchmarking Universal Single-Copy Orthologs (BUSCO) (Simão *et al.* 2015) (using the ‘Nematoda_ob9’ dataset). For each species, the highest quality assembly was selected and, where possible, scaffolded with assembled transcripts using SCUBAT2 (https://github.com/GDKO/SCUBAT2) or with long-insert “mate-pair” data using Platanus.

We identified repeats independently in each genome using RepeatModeler (Smit & Hubley 2010). After filtering sequences that likely originated from protein-coding genes, we combined each repeat library with known Rhabditida repeats obtained from RepBase (Jurka *et al.* 2005). This concatenated repeat library was then provided to RepeatMasker (Smit *et al.* 1996) for masking. If RNA-seq data were available, reads were aligned to the assembly using STAR (Dobin *et al.* 2013) and the resulting BAM file provided to BRAKER (Hoff *et al.* 2016), which performed final gene prediction. If RNA-seq data were not available, genes were predicted initially using MAKER2 (Hoff *et al.* 2016) with a training set composed of *ab initio* predictions from GeneMark (Lukashin & Borodovsky 1998), gene models identified by CEGMA (Parra *et al.* 2007), and the *C. elegans* protein sequence set. The resulting gene models were used as to train AUGUSTUS (Keller *et al.* 2011), which generated the final gene set.

### Phylogenetic analysis

Accessions to all data used in phylogenomics analysis are available in Table S6. For those species with available genome sequences, we identified and collected the protein sequence of the longest isoform of each protein-coding gene. For those species for which only transcriptome data was available (*C. guadeloupensis* and *C. vivipara*), open reading frames and putative protein sequences were predicted using TransDecoder (Haas & Papanicolaou 2012). OrthoFinder (Emms & Kelly 2015) (using the default inflation value of 1.5) was used to cluster all protein sequences into putatively orthologous groups (OGs). OGs containing loci present as single-copy in at least 27 of the 33 species (except in those species with genome assemblies known to contain regions of uncollapsed heterozygosity [i.e., *C. angaria, C. brenneri, C. japonica, C. waitukubuli* and *D. coronatus*] and in those species for which only transcriptome data were available, where counts of two were allowed) were selected.

To identify paralogous sequences, we aligned the protein sequences of each OG using FSA (Bradley *et al.* 2009) and generated a maximum likelihood tree along with 100 rapid bootstraps using RAxML (Stamatakis 2014)(using best-fitting amino acid substitution, as selected by the model testing component of RAxML, and gamma-distributed rate-variation among sites). Each tree was screened by PhyloTreePruner (Kocot *et al.* 2013)(collapsing nodes with bootstrap support < 50), and any OGs containing paralogues were discarded. If two representative sequences were present for any species (i.e., “in-paralogues”) after this paralogue screening step, the longest of the two sequences was retained and the other discarded.

The protein sequences of each one-to-one OG were then aligned using FSA and gene trees estimated as previously described. ASTRAL-III (Mirarab & Warnow 2015), a coalescent-based method, was then used to reconstruct the species phylogeny using individual genes trees as an input. We also reconstructed the species tree using a concatenation approach. TrimAl (Capella-Gutiérrez *et al.* 2009) was used to remove spuriously aligned regions from each alignment, which were subsequently concatenated using catfasta2phyml (available at https://github.com/nylander/catfasta2phyml). Maximum likelihood (ML) analysis was performed with RAxML (general-time reversible model (GTR) (Tavaré 1986) with gamma-distributed rate-variation among sites) along with 100 bootstrap replicates. Bayesian inference was carried out using the site-heterogeneous CAT-GTR substitution model (Lartillot & Philippe 2004) (with gamma-distributed rate-variation among sites) implemented in PhyloBayes MPI (Lartillot *et al.* 2013), with two independent chains. Convergence was assessed using Tracer (Rambaut *et al.* 2007). A posterior consensus tree was estimated using samples from both chains, with the initial 10% of all trees discarded as burn-in. Newick trees were visualised using the iTOL web server (Letunic & Bork 2016).

### Gene structure analysis

For those species for which gene structure information was available, we collected the longest isoform of each protein-coding gene. As previously described, we clustered protein sequences into putatively orthologous groups using OrthoFinder, and removed redundant duplicate sequences and discarded clusters containing paralogues. We identified OGs which were present as single-copy in all 28 species. We extracted gene structure information using the Ensembl Perl API (Zerbino *et al.* 2018) and calculated total intron count each species. Correlations between genome size and intron count were calculated using Pearson’s correlation coefficient (r), implemented in R (Team & Others 2013).

### Notch-like Receptor Analysis

In an existing orthology clustering set, we identified the OG that contained the *C. elegans* proteins LIN-12 (R107.8) and GLP-1 (F02A9.6) and collected the protein sequences of each member. After removing sequences that were shorter than 700 amino acids, we generated an amino acid alignment using FSA. We performed a maximum likelihood analysis using RAxML, allowing the substitution model to be automatically selected, and conducted 100 rapid bootstrap replicates. Branch lengths were extracted using a custom Python script (available at https://github.com/lstevens17/caeno-ten-descriptions), making use of the ete3 module (Huerta-Cepas *et al.* 2016). We identified conserved domains in each protein sequence using InterProScan (Jones *et al.* 2014). Counts of EGF-repeats were obtained from were obtained from the ProSiteProfiles database (release 2017_09)(Sigrist *et al.* 2013).

### Scanning electron microscopy

Nematode cultures were resuspended and washed twice in M9 solution, then fixed overnight at 4°C in M9 or 50 mM phosphate pH 7.0 + glutaraldehyde 2.5 to 4%, depending on the batch. The fixed animals were rinsed twice in M9 and dehydrated through an ethanol series, pelleting them at each step at 1 g in a tube. The samples were processed through critical point drying and coating with 20 nm of Au/Pd, and then observed with a JEOL 6700F microscope at the Ultrastructural Microscopy Platform of the Pasteur Institute.

### Nomarski micrographs

The Nomarski micrographs were taken using an AxioImager 2 (Zeiss) after mounting the animals on a noble agar pad as described in (Shaham 2006). The pictures showing extruded spicules were taken after exposing the animals for 2 seconds in the microwave before adding the coverslip.

## Acknowledgements

We dedicate this paper, and *Caenorhabditis sulstoni*, to John Sulston, a mentor and colleague. We thank Adeline Mallet for assistance at the Ultrastructural Microscopy Platform of the Pasteur Institute. We are very grateful to Fabrice Besnard, Ludmilla Lokmane, Jean-Baptiste Pénigault, Clotilde Gimond, Charlie Gosse, Howard Baylis, Kevin Howan, Sarah Mühlberger, Amir Yassin for samples. We thank Taisei Kikuchi, Janna Fierst and Erich Schwarz for pre-publication access to the access to genomic data. We gratefully acknowledge the Republic of Panama and the Smithsonian Tropical Research Institute for supporting collections on Barro Colorado Island. Samples were exported from Panama under Scientific Permit SEX/A-25-12 from the Autoridad Nacional del Ambiente. We thank the staff of Edinburgh genomics for expert support. Edinburgh Genomics has core funding from the UK Natural Environment Research Council (UKSBS PR18037). LS was supported by a Baillie Gifford studentship. LF, AR and MAF were supported by the Centre National de la Recherche Scientifique (CNRS), the Ecole Normale Supérieure and by Agence Nationale pour la Recherche grant ANR-11-BSV3-013. CB and SF were supported by the CNRS. KCK and DHAF were supported by NSF grant NSF IOS-1656736. MVR was supported by NIH grant R01GM121828.

## Author Contributions

CB, MAF, LF, TK, MVR, and WS collected and isolated nematodes. CB, LF, SF, MAF, MVR and TK generated inbred lines. CB, MAF, MVR, and LF performed mating tests. CB, SF, LN and AR isolated DNA and RNA. TB, CC, MDN, LN and LS performed genome assembly and annotation. MAF, DHAF, KK, and WS studied morphology. LS performed phylogenomic and comparative genomic analysis. LS, MAF and MB wrote the manuscript. All authors read, provided comments on and approved the final version of the manuscript.

## Data Accessibility

All raw sequence data, reference genome assemblies and rDNA sequences have been deposited in the relevant INSDC databases (see Table 1 for accession numbers). Accessions and links to all data used in the phylogenomics analysis are available in Table S6. Data files associated with this study have been deposited in Zenodo under the accession 10.5281/zenodo.1402254. All assemblies and annotations described in this paper are are available to browse, query, and download at caenorhabditis.org. Information about morphology, geographic distribution, selected gene sequence data and taxonomy of these and other *Caenorhabditis* species has been deposited in RhabditinaDB (rhabditina.org).

## Supporting Information

**Table S1: List of isolates and their origin.**

**Table S2: Mating tests.** This table contains several sheets, showing the results of crosses between isolates of different species.

**Table S3: Detailed genome assembly and gene prediction statistics.**

**Table S4: Mean branch lengths from Maximum likelihood gene tree of all Notch-like proteins.** Branch lengths were extracted using a custom Python script (available at https://github.com/lstevens17/caeno-ten-descriptions)

**Table S5: EGF-repeat counts for LIN-12/GLP-1 homologues.** Counts of EGF-repeats were obtained from were obtained from the ProSiteProfiles database (release 2017_09).

**Table S6: Accessions and links to data used in phylogenomic analysis.**

**Figure S1: Kmer spectra for *C. waitukubuli* (sp. 39).** Kmers were counted using KMC (v2.3). Plotted in R using the ggplot2 package.

**Figure S2: PhyloBayes phylogenetic tree.** Phylogenetic tree inferred using Bayesian inference with the CAT-GTR+Γ substitution model. Bayesian posterior probabilities are 1.0 unless noted as branch annotations. Scale is in substitutions per site.

**Figure S3: RAxML phylogenetic tree.** Maximum likelihood phylogenetic inferred using RAxML with the GTR+Γ substitution model. Bootstrap support values (100 replicates) are 100 unless noted as branch annotations. Scale in substitutions per site.

**Figure S4: ASTRAL-III phylogenetic tree.** Phylogenetic tree inferred using ASTRAL-III, by providing maximum likelihood gene trees (inferred using RAxML with the substitution model selected automatically) as input. As ASTRAL-III outputs trees with branch lengths in coalescent units, branch lengths in substitutions per site were estimated using RAxML with the GTR+Γ substitution model and the concatenated alignment. Bayesian posterior probabilities are 1.0 unless noted as branch annotations. Scale is in substitutions per site.

**Figure S5: Morphology of *C. parvicauda* male tail.** (A, A’) Two focal planes in Nomarski optics of the same adult male tail of strain NIC134. (B) Scanning electron microscopy, strain JU2070. ad: anterior dorsal papilla; pd: posterior dorsal papilla; vr1, etc.: ventral papilla 1, etc. These photographs exemplify the left-right asymmetry of the posterior papillae, observed in strains NIC134 and JU2070, as well as JU1766 (not shown).

**Figure S6: Morphology of *C. quiockensis* and *C. vivipara*.** (A-C) Nomarski micrographs of male tails of *C. quiockensis* strain JU2745. D. *C. castelli* JU1426 is shown for comparison. Same scale in A-D. (E) C. vivipara NIC1070 male tail, with details of pericloacal regions of other animals in (F-H). Note the much larger and wider fan in *C. vivipara*. Ad: anterior dorsal papilla; gub: gubernaculum; pd: posterior dorsal papilla; spic: spicule; v1, etc.: ventral papillae.(A,B,D, H) are lateral views. (C) and (E-G) are ventral views. I. Gravid female adult with late-stage embryos in the uterus and a recently laid L1 stage larva. Bars: 10 µm, except 5 µm for (F,G) and 50 µm for (I). Anterior is to the left except in H where the anterior is to the top.

**Figure S7: Male tails of species of the Elegans supergroup.** Ventral views by Nomarski optics of *C. becei* strain QG704, *C. waitukubuli* strain NIC564, *C. panamensis* strain QG702, *C. sulstoni* strain SB454 and *C. afra* JU1199 male tails. Note the variation in the respective lengths of rays 4 and 5. All pictures at are at the same scale. Bar: 10 µm. The correspondence between the v1-7, ad, pd papilla nomenclature and that of rays used in *C. elegans* is indicated on the *C. becei* picture. v1-7 denote ventral papillae, while ad and pd denote the anterior and posterior dorsal papillae, respectively. In the *C. elegans* nomenclature, rays are instead numbered r1-r9 without distinction between ventral and dorsal rays.

**Figure S8: Spicule tip shape in Elegans supergroup species.** Nomarski micrographs. Left column: ventral view. Right column: ventro-lateral view. The tip of spicules in *C. afra* and *C. sulstoni* (as in several species of the Japonica group; (Kiontke et al. 2011) is broad and bent with a discontinuity on the curvature, compared to a thin and continuously bent tip in species of the Elegans group such as *C. zanzibari* or *C. elegans*. That of *C. panamensis* strain QG702 is quite thin, while that of *C. becei* strain QG704 is broad. Bar for all panels: 5 µm.

**Document S1: Species declarations.**

**Document S2: Detailed bioinformatics methods.** Methods, versions and relevant parameters used in genome assembly, gene prediction, phylogenomic analysis, Notch-receptor analysis and gene structure analysis.

